# Genome-wide investigation of multiplexed CRISPR-Cas12a-mediated editing in rice

**DOI:** 10.1101/2022.07.31.502228

**Authors:** Yingxiao Zhang, Yuechao Wu, Gen Li, Aileen Qi, Yong Zhang, Tao Zhang, Yiping Qi

## Abstract

We previously reported highly specific genome editing in rice by CRISPR-Cas9 and Cas12a with single DNA double strand break (DSB) (Tang et al., 2018). Two concurrent DSBs by CRISPR-Cas9 could generate defined deletions (Zhou et al., 2014), inversions (Schmidt et al., 2020), and translocations (Beying et al., 2020). Off-target effects of many simultaneous DSBs are unknown in plants. Here, we used whole-genome sequencing (WGS) to investigate off-target effects in rice plants edited by highly multiplexable CRISPR-Cas12a systems (Zhang et al., 2021). Our WGS study revealed highly specific multiplexed genome editing by CRISPR-Cas12a. We found low-frequency large chromosomal deletion and duplication events in plants that endured many (e.g., >50) simultaneous DSBs, but not in plants that endured a lower order DSBs (e.g., <10). While our short reads sequencing may not capture all chromosomal rearrangements, our results nevertheless shed important light on the analysis and regulation of engineered crops derived from multiplexed genome editing.

---

We previously reported highly specific genome editing in rice by CRISPR-Cas9 and Cas12a with single DNA double strand break (DSB) (Tang et al., 2018). Two concurrent DSBs by CRISPR-Cas9 could generate defined deletions (Zhou et al., 2014), inversions (Schmidt et al., 2020), and translocations (Beying et al., 2020). Off-target effects of many simultaneous DSBs are unknown in plants. Here, we used whole-genome sequencing (WGS) to investigate off-target effects in rice plants edited by highly multiplexable CRISPR-Cas12a systems (Zhang et al., 2021).

In the first scenario, we pursued lower order multiplexed editing by targeting four sites (T1 to T4) with the canonical TTTV protospacer adjacent motif (PAM) in the rice genome using either LbCas12a or Mb2Cas12a (Figure 1A). In the second scenario, a notably higher order LbCas12a-mediated multiplexed editing was conducted: 12 additional CRISPR RNAs (crRNAs) were co-expressed with the previous four crRNAs to target 16 canonical TTTV PAM sites (T5-TTTV to T16-TTTV). By our design, 11 of the 12 crRNAs (T6 to T16) could potentially each target a second site with relaxed TTV PAMs. If LbCas12a could edit TTV PAMs as FnCas12a (Zhong et al., 2018) and Mb2Cas12a (Zhang et al., 2021) do, this higher order construct would edit up to 27 target sites across 11 chromosomes, generating >50 concurrent DSBs per diploid cell (Figure 1A).

**Figure 1.**
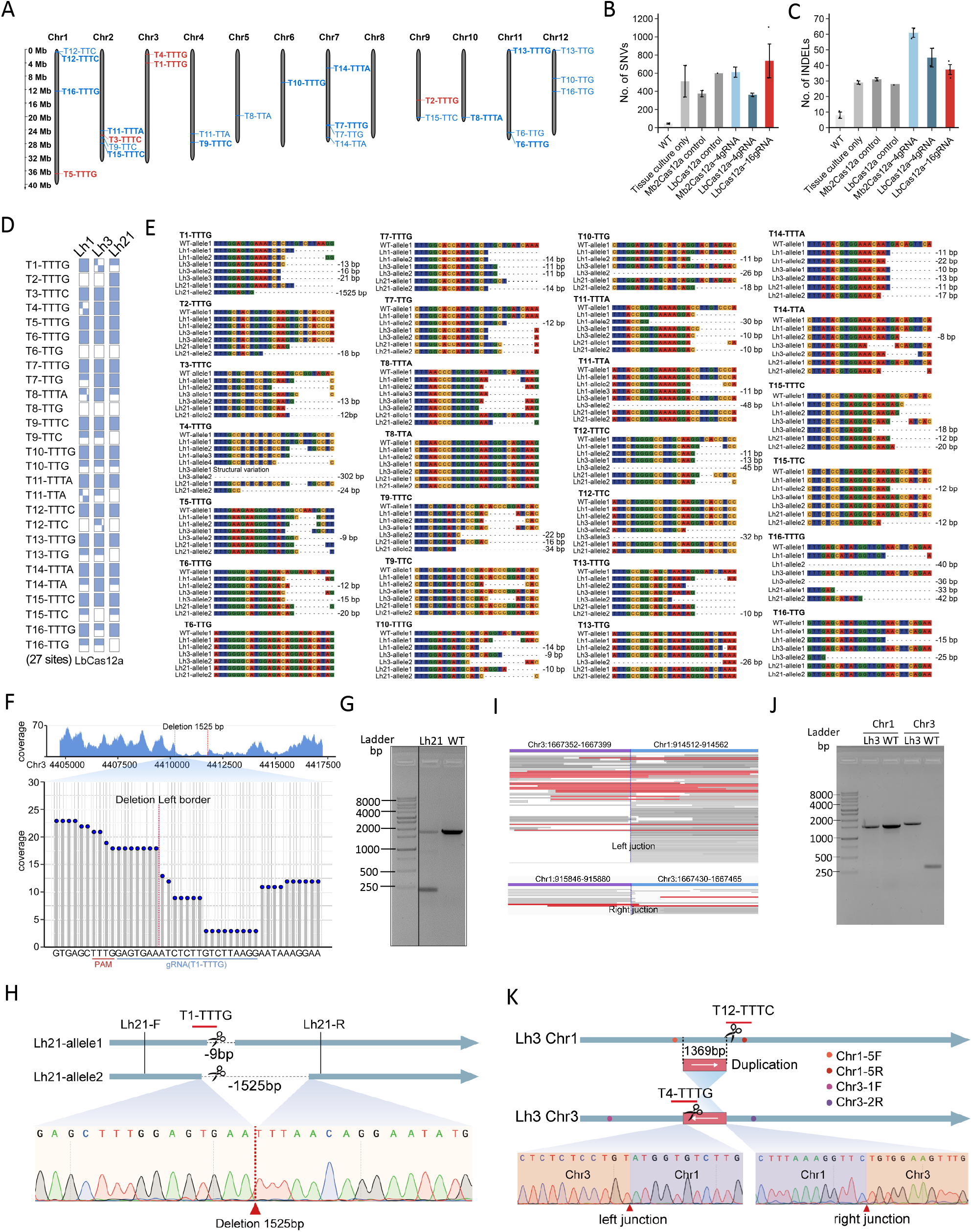
WGS-based on- and off-target analyses of multiplexed Cas12a genome editing in rice. (A) Schematics of simultaneous editing at 27 target sites across rice chromosomes through multiplex 16 crRNAs, with TTTV PAM (bold) and TTV PAM (nonbold), one editing site (red) and two editing sites (blue). (B, C) Numbers of whole genome SNVs (B) and INDELs (C) per plant, not counting on-target INDELs. Error bars represent standard error of the mean (SEM) of two or three lines. (D) Zygosity analysis of three LbCas12a T0 lines (Lh1, Lh3 and Lh21) at 27 target sites. Wild type (empty rectangle), monoallelic mutant (half-filled rectangle), biallelic mutant (fully filled rectangle) and chimeric mutant (gridded rectangle) are indicated for each line and target sites. (E) Genotypes of three LbCas12a T0 at 27 target sites. (F) Top panel, the reads coverage of the 5kb flanking region of the deletion. Bottom panel, the reads distribution of the target site T1-TTTG. (G) Validation of large deletion in Lh21 by gel electrophoresis of amplicons (using primers Lh21-F/R denoted in H). (H) Validation of 1525 bp deletion in Lh21 by Sanger sequencing. (I) IGV browser views showing the reads coverages at junction of translocation, reads across the junctions are indicated in red. (J) Validation of duplication of Chr1 fragment into Chr3 in Lh3 by gel electrophoresis of amplicons (using primers Chr1-5F/5R and Chr3-1F/2R denoted in K). (K) Validation of duplicated fragment from Chr1 in Chr3 by Sanger sequencing. Duplicated fragment is displayed as red bar with arrow.

Three multiplexed constructs (Supplemental Figure 1-3) and their no-crRNA controls were used for rice transformation. T0 lines carrying high levels of multiplexed editing were selected for WGS, including two LbCas12a lines containing four crRNAs (L30 and L32), two Mb2Cas12a lines containing four crRNAs (M9 and M10), and three LbCas12a lines containing 16 crRNAs (Lh1, Lh3 and Lh21). The WGS data were analyzed to identify single nucleotide variations (SNVs) and insertions/deletions (INDELs) (Supplemental Figure 4 and Supplemental Table 1) (Tang et al., 2018). The multiplexed genome editing constructs generated ~400 to ~700 SNVs, which are comparable to the controls (Figure 1B). The Mb2Cas12a construct generated twice as many INDELs (~60) as controls (~30) (Figure 1C). However, it shared a similar deletion length pattern to other constructs (Supplemental Figure 5). Furthermore, among all constructs, no off-target mutations were detected at putative off-target sites (Supplemental Table 2). By contrast, all four lines (L30, L32, M9 and M10) generated biallelic on-target editing with variable lengths of small deletions (Supplemental Figure 6). These data suggest MbCas12a has high targeting specificity.

For the higher order LbCas12a-mediated editing using 16 crRNAs, 21 T0 lines were used for genotype analysis using NGS of PCR amplicons. The editing efficiencies at canonical TTTV PAM sites were higher than at the TTV sites (Supplemental Table 4). We focused our analysis on the three T0 lines (Lh1, Lh3 and Lh21) that simultaneously express 16 crRNAs with LbCas12a. WGS revealed that 15 out 16 TTTV target sites were edited in Lh1 and Lh3. Moreover, all 16 TTTV target sites were edited in Lh21 (Figure 1D), demonstrating highly efficient multiplexed Cas12a genome editing as we recently reported (Zhang et al., 2021). Strikingly, many target sites with non-canonical TTV PAMs were also edited by LbCas12a. Out of the 11 TTV PAM sites, eight in Lh1, nine in Lh3, and four in Lh21 have been edited with ~10-40 bp deletions, respectively (Figure 1D and 1E). The canonical TTTV PAM sites are preferred by LbCas12a with higher editing efficiencies over the TTV PAM sites (Supplemental Figure 7). This finding demonstrates LbCas12a can edit many TTV PAMs albeit with lower efficiency.

We further investigated the chromosome rearrangements in multiplex edited rice lines. Among all four T0 lines that carry four biallelic mutations by LbCas12a and Mb2Cas12a, no evidence of chromosomal rearrangements was found. For the T0 lines multiplexing 16 crRNAs, no chromosomal rearrangement was detected in Lh1. However, we identified a large deletion in Lh21 (Figure 1F). We confirmed this large deletion of 1525 bp using PCR (Figure 1G) and Sanger sequencing (Figure 1H). Interestingly, the deletion started at the T1-TTTG target site, and the other end was not defined by any target site (Figure 1H). Since 20 out of 27 target sites were edited in Lh21, a heavy DNA repair burden might have caused the large deletion.

In line Lh3, WGS revealed that a segment of Chromosome 1 near the T12-TTTC target site was translocated onto Chromosome 3 at the T4-TTTG target site (Figure 1I). PCR and Sanger sequencing showed this translocation event of being 1369 bp in length (Figure 1J, 1K, and Supplemental Figure 8). Our further analysis showed this fragment was not deleted on Chromosome 1 (Figure 1J and Supplemental Figure 9), indicating the event is a duplication. There is a 4 bp microhomology sequence between the two target sites on Chromosome 1 and Chromosome 3. Hence, we proposed a plausible microhomology based recombination model like synthesis-dependent strand annealing (SDSA) (Puchta, 2005) to explain this duplication event (Supplemental Figure 10).

In summary, our WGS study revealed highly specific multiplexed genome editing by CRISPR-Cas12a. We found low-frequency large chromosomal deletion and duplication events in plants that endured many (e.g., >50) simultaneous DSBs, but not in plants that endured a lower order DSBs (e.g., <10). While our short reads sequencing may not capture all chromosomal rearrangements, our results nevertheless shed important light on the analysis and regulation of engineered crops derived from multiplexed genome editing.

## ACKNOWLEDGEMENTS

This work was supported by the USDA-NIFA BRAG program (award no. 2020-33522-32274) to Y. Q.

## COMPETING INTERESTS

The authors declare no competing financial interests.

## AUTHOR CONTRIBUTIONS

Y.Q. and Yingxiao.Z. designed the study. Y.Q., Yong.Z. and T.Z. supervised the study. Yingxiao Z., G.L. and A. Q. performed the experiments and analyzed the data. Y.W. and T.Z. analyzed the whole genome sequencing data. Y.W., Yingxiao.Z. and G.L. made the figures. Y.Q. wrote the first draft of the manuscript. All authors revised and approved the final version.

## Notes

### Competing Interest Statement

The authors have declared no competing interest.

