## Supplemental Methods, Tables and Figures for "Genome-wide investigation of multiplexed CRISPR-Cas12a-mediated editing in rice"

[1]Department of Plant Science and Landscape Architecture, University of Maryland, College Park, Maryland 20742, USA; [2]Jiangsu Key Laboratory of Crop Genomics and Molecular Breeding / Jiangsu Key Laboratory of Crop Genetics and Physiology , Agricultural College of Yangzhou University, Yangzhou 225009, China; [3]Jiangsu Co-Innovation Center for Modern Production Technology of Grain Crops, Yangzhou University, Yangzhou 225009, China; [4]Department of Biotechnology, School of Life Science and Technology, Center for Informational Biology, University of Electronic Science and Technology of China, Chengdu 610054, China; [5]Key Laboratory of Plant Functional Genomics of the Ministry of Education/Joint International Research Laboratory of Agriculture and Agri-Product Safety, The Ministry of Education of China, Yangzhou University, Yangzhou 225009, China; [6]Institute for Bioscience and Biotechnology Research, University of Maryland, Rockville, Maryland 20850, USA.

[†] These authors contributed equally to this work.

[#] Present address: Syngenta, Research Triangle Park, North Carolina 27709, USA.

***Corresponding authors**

Yiping Qi, Department of Plant Science and Landscape Architecture, University of Maryland, College Park, MD 20742, USA;

Tao Zhang, Agricultural College of Yangzhou University, Yangzhou 225009, China;

- Supplemental Materials and Methods.

- Supplemental Table 1. Summary of sequencing depth and coverage of WGS samples.
- Supplemental Table 2. Summary of off-target mutations at potential off-target sites in rice genome.
- Supplemental Table 3. Oligos used in this study.
- Supplemental Table 4. Genome editing efficiency in T0 lines by LbCas12a with multiplexing 16 crRNAs.

- Supplemental Figure 1. Vector map for LbCas12a based simultaneous expression of four crRNAs in the rice genome.
- Supplemental Figure 2. Vector map for Mb2Cas12a based simultaneous expression of four crRNAs in the rice genome.
- Supplemental Figure 3. Vector map for LbCas12a based simultaneous expression of 16 crRNAs in the rice genome.
- Supplemental Figure 4. Whole genome sequencing and analysis pipeline.
- Supplemental Figure 5. Deletion size analysis of WGS samples.
- Supplemental Figure 6. Genotypes of T0 lines with multiplexing four crRNAs by Mb2Cas12a and LbCas12a.
- Supplemental Figure 7. Averaged editing efficiency at different target sites in T0 lines by LbCas12a with multiplexing 16 crRNAs.
- Supplemental Figure 8. PCR validation of a large DNA insertion on Chromosome 3.
- Supplemental Figure 9. Duplicated DNA sequence in Chromosome 3 is not deleted in Chromosome 1.
- Supplemental Figure 10. Microhomology-mediated synthesis-dependent strand annealing involved in duplication of Chr1 fragment in Chr3.

### Supplemental Materials and Methods

#### Plant material and generation of mutated lines

*Oryza sativa* L.ssp. Japonica cv. Nipponbare was used in this study. Rice mutated lines were generated based on highly efficient multiplexed CRISPR-Cas12 systems described in a previous study (Zhang et al., 2021). The T-DNA vector pLR816 was transformed into rice to generate LbCas12a T0 lines containing 4 crRNAs. The T-DNA vector pLR1766 was transformed into rice to generate Mb2Cas12a T0 lines containing 4 crRNAs. The T-DNA vector pLR1963 was transformed into rice to generate LbCas12a T0 lines containing 16 crRNAs. These three transformations were performed from different explant batches derived from the same seeds. Regularly grown WT plants and tissue culture derived plants from one batch were used as controls. Leaf tissue of each control and T0 lines was collected for DNA extraction using the CTAB method.

#### Whole genome sequencing

Genomic DNA was extracted from leaf tissue of T0 plants using DNeasy® Plant Mini Kit (QIAGEN, Germany). DNA samples were submitted to Genewiz for whole genome sequencing according to the submission guideline. Illumina sequencing was performed by Genewiz on a HiSeq 4000 platform (Genewiz, USA). The WGS raw data reported in this article have been deposited to the Sequence Read Archive in National Center for Biotechnology Information (NCBI) under the accession numbers PRJNA718633 and Beijing Institute of Genomics Data Center (http://bigd.big.ac.cn) under BioProject PRJCA007567.

#### Whole genome sequencing analysis

WGS analysis was conducted following our previous research (Tang et al., 2018). Briefly, the reads adapter was trimmed by applying SKEWER (v. 0.2.2) (Jiang et al., 2014). Clean high quality reads were mapped to rice (cv. Nipponbare) reference genome TIGR7(http://rice.uga.edu/) using BWA mem (v. 0.7.17) (Li and Durbin, 2010). Picard and Samtools (v. 1.9)(Li et al., 2009) were used to filter multiple mapping reads and generate sorted BAM files. Reads near indels were realigned using GATK (v. 3.8) (McKenna et al., 2010) . SNVs and INDELS across the entire gnome were detected using LoFreq (v. 2.1.2) (Wilm et al., 2012), Mutect2 (Cibulskis et al., 2013), VarScan2(v. 2.4.3) (Koboldt et al., 2012) and Pindel(v.0.2.5)(Chen et al., 2016). Bedtools (v. 2.27.1) (Li, 2011) were used overlapping SNVs and indels. Manta (v.1.6.0) (Chen et al., 2016) was used to detect structural variation and large mutations. The potential off-target sites were predicted by Cas-OFFinder (v. 2.4) (Bae et al., 2014) with up to 5-nt mismatches, PAM is set to TTV. Data processing and analysis were done using python and R.

**Mutation analysis by PCR and Sanger sequencing**

To verify mutation variants, PCR was performed using Q5® High-Fidelity DNA Polymerase (New England Biolabs, USA). 5 µl PCR products for each sample were visualized on 1% agarose gel. PCR products were gel-purified or enzymatically purified by Exonuclease I and quick CIP (New England Biolabs, USA) with incubation at 37°C for 30 min followed by at 80°C for 20 min. Purified PCR products were used for Sanger sequencing (Genewiz, USA).

### Supplemental Table 1. Summary of sequencing depth and coverage of WGS samples.

| group | WGS ID | Sequencing depth(X) | mapping ratio (%) | Genome coverage (%) |
|---|---|---|---|---|
| WT | W1 | 52.43 | 93.97 | 99.73 |
| WT | W2 | 53.21 | 94.25 | 99.73 |
| Tissue culture only | T1 | 46.84 | 93.36 | 99.35 |
| Tissue culture only | T2 | 50.24 | 94.9 | 99.13 |
| Mb2Cas12a backbone control | Mc3 | 55.84 | 93.85 | 99.38 |
| Mb2Cas12a backbone control | Mc4 | 63.49 | 93.27 | 99.61 |
| LbCas12a backbone control | Lc3 | 62.87 | 93.81 | 99.67 |
| Mb2Cas12a-4gRNA | M9 | 54.38 | 94.38 | 99.59 |
| Mb2Cas12a-4gRNA | M10 | 46.62 | 94.16 | 99.69 |
| LbCas12a-4gRNA | L30 | 59.15 | 93.94 | 99.81 |
| LbCas12a-4gRNA | L32 | 62.74 | 93.61 | 99.8 |
| LbCas12a-16gRNA | Lh1 | 43.07 | 88.67 | 99.68 |
| LbCas12a-16gRNA | Lh3 | 49.28 | 88.18 | 94.88 |
| LbCas12a-16gRNA | Lh21 | 52.15 | 91.21 | 99.67 |

**Supplemental Table 2. Summary of off-target mutations at potential off-target sites in rice genome.**

| Nucleotide mismatch | Line | LbCas12a-4crRNAs | Mb2Cas12a-4crRNAs | LbCas12a-16crRNAs |
|---|---|---|---|---|
| 1 | T0 plant 1 | 0/0 | 0/0 | 0/0 |
| | T0 plant 2 | 0/0 | 0/0 | 0/0 |
| | T0 plant 3 | -- | -- | 0/0 |
| ⩽2 | T0 plant 1 | 0/0 | 0/0 | 0/1 |
| | T0 plant 2 | 0/0 | 0/0 | 0/1 |
| | T0 plant 3 | -- | -- | 0/1 |
| ⩽3 | T0 plant 1 | 0/0 | 0/0 | 0/7 |
| | T0 plant 2 | 0/0 | 0/0 | 0/7 |
| | T0 plant 3 | -- | -- | 0/7 |
| ⩽4 | T0 plant 1 | 0/4 | 0/4 | 0/20 |
| | T0 plant 2 | 0/4 | 0/4 | 0/20 |
| | T0 plant 3 | -- | -- | 0/20 |
| ⩽5 | T0 plant 1 | 0/35 | 0/35 | 0/102 |
| | T0 plant 2 | 0/35 | 0/35 | 0/102 |
| | T0 plant 3 | -- | -- | 0/102 |

**Supplemental Table 3. Oligos used in this study.**

| Oligo name | Oligo sequence 5'-3' |
|---|---|
| Chr1-5F | CGGTTAATAGTGCGGAAAACATG |
| Chr1-5R | CATGGAGACGAGGAGTACACAAG |
| Chr1-F1 | ACGTCTTAGCCATTAACGCA |
| Chr1-R1 | GCAGGAGCCCAGCTGTATAA |
| Chr1-F2 | CTTGGACATGGACGACGTGA |
| Chr1-R2 | AGTAGGCTGGACCGGGTATT |
| Chr1-F3 | TTGATCCGAATGGTGGCCTT |
| Chr1-R3 | CGGCTCATGCAGAATGTTCG |
| Chr1-F4 | TTGACAAAGCCGACCCAAGT |
| Chr1-R4 | GCAACCTAGGACCAGATGTGA |
| Chr1-F5 | GCAAATCCACCCCTCATGGA |
| Chr1-R5 | ATGACTTAATGTGCGAGAAAGC |
| Chr3-1F | ACCATGGCCGGGGTACAAATA |
| Chr3-2R | GAAAAGTTTCTCCTACCGCTGC |
| Chr3-R1 | ACAACGGCATACGCCCAAAT |
| Lh21-F | TGGAACCAGCCAAGCAAGAT |
| Lh21-R | GCTGGCAAAAGTCCAAGAGC |
| OsPDS-HTS-R2 | GCTCATGATATTTATGTGACGTTAA |

**Supplemental Table 4. Genome editing efficiency in T0 lines by LbCas12a with multiplexing 16 crRNAs.**

| Target sites | Gene ID | crRNA sequence 5'-3' | Editing efficiency (%)* | Biallilic editing efficiency (%)* |
|---|---|---|---|---|
| T1-TTTG | Os03g08570 (OsPDS) | GAGTGAAATCTCTTGTCTTAAGG | 95.2 | 90.5 |
| T2-TTTG | Os09g26999 (OsDEP1) | CTACTGTTGCAAGTGCTCACCCA | 14.3 | 5.8 |
| T3-TTTC | Os02g45250 (OsROC5) | TGCTTCCTGCAATGCCGGTAGAC | 95.2 | 61.9 |
| T4-TTTG | Os03g03724 (OsmiR528) | CCTCTCTCTCCTGTGCTTGCCTC | 76.2 | 47.6 |
| T5-TTTG | Os01g68598 (OsEPFL9) | AAGAAGGGTTATGGCCAATGCTT | 100.0 | 100.0 |
| T6-TTTG | Os11g44430 | GGGCATGGAGACAGGAGACATAG | 100.0 | 81.0 |
| T6-TTG | Os11g44260 | GGGCATGGAGACAGGAGACATAG | 4.8 | 0.0 |
| T7-TTTG | Os07g40404 | GCACCATATGCTTGCTGATCAAA | 100.0 | 95.2 |
| T7-TTG | Os07g40790 | GCACCATATGCTTGCTGATCAAA | 100.0 | 100.0 |
| T8-TTTA | Os10g40824 | ACCCTGTGTGAATGGTCAGTAAG | 71.4 | 5.8 |
| T8-TTG | Os05g36070 | ACCCTGTGTGAATGGTCAGTAAG | 100.0 | 14.3 |
| T9-TTTC | Os04g50120 | TGTATCTCCGACACCCGGATCAC | 95.2 | 71.4 |
| T9-TTC | Os02g46610 | TGTATCTCCGACACCCGGATCAC | 28.6 | 0.0 |
| T10-TTTG | Os06g18810 | GATGATGCATCAGGTACTAGAAC | 90.5 | 52.4 |
| T10-TTG | Os12g16490 | GATGATGCATCAGGTACTAGAAC | 95.2 | 4.8 |
| T11-TTTA | Os02g43194 | CCGGTGAAAAGGACCTTGTCCCA | 100.0 | 76.2 |
| T11-TTA | Os04g45720 | CCGGTGAAAAGGACCTTGTCCCA | 33.3 | 0.0 |
| T12-TTTC | Os01g02690 | TGGGGCCTTGCAAGGTCACCTCC | 95.2 | 52.4 |
| T12-TTC | Os01g02420 | TGGGGCCTTGCAAGGTCACCTCC | 19.0 | 0.0 |
| T13-TTTG | Os11g01450 | CCGGCAGCTAATAGGGATCTAAA | 95.2 | 81.0 |
| T13-TTG | Os12g01480 | CCGGCAGCTAATAGGGATCTAAA | 95.2 | 0.0 |
| T14-TTTA | Os07g10860 | TACGTGGAAACAATGACAGTTCA | 100.0 | 100.0 |
| T14-TTA | Os07g47284 | TACGTGGAAACAATGACAGTTCA | 0.0 | 0.0 |
| T15-TTTC | Os02g49270 | TCCTGAGGAGCAAGAGCCATCAC | 100.0 | 100.0 |
| T15-TTC | Os09g37860 | TCCTGAGGAGCAAGAGCCATCAC | 0.0 | 0.0 |
| T16-TTTG | Os01g23900 | AGCATATGGTTGTAACTTCAGAA | 100.0 | 95.2 |
| T16-TTG | Os12g24050 | AGCATATGGTTGTAACTTCAGAA | 28.6 | 0.0 |

*Editing efficiency was calculated based on 21 T0 lines.

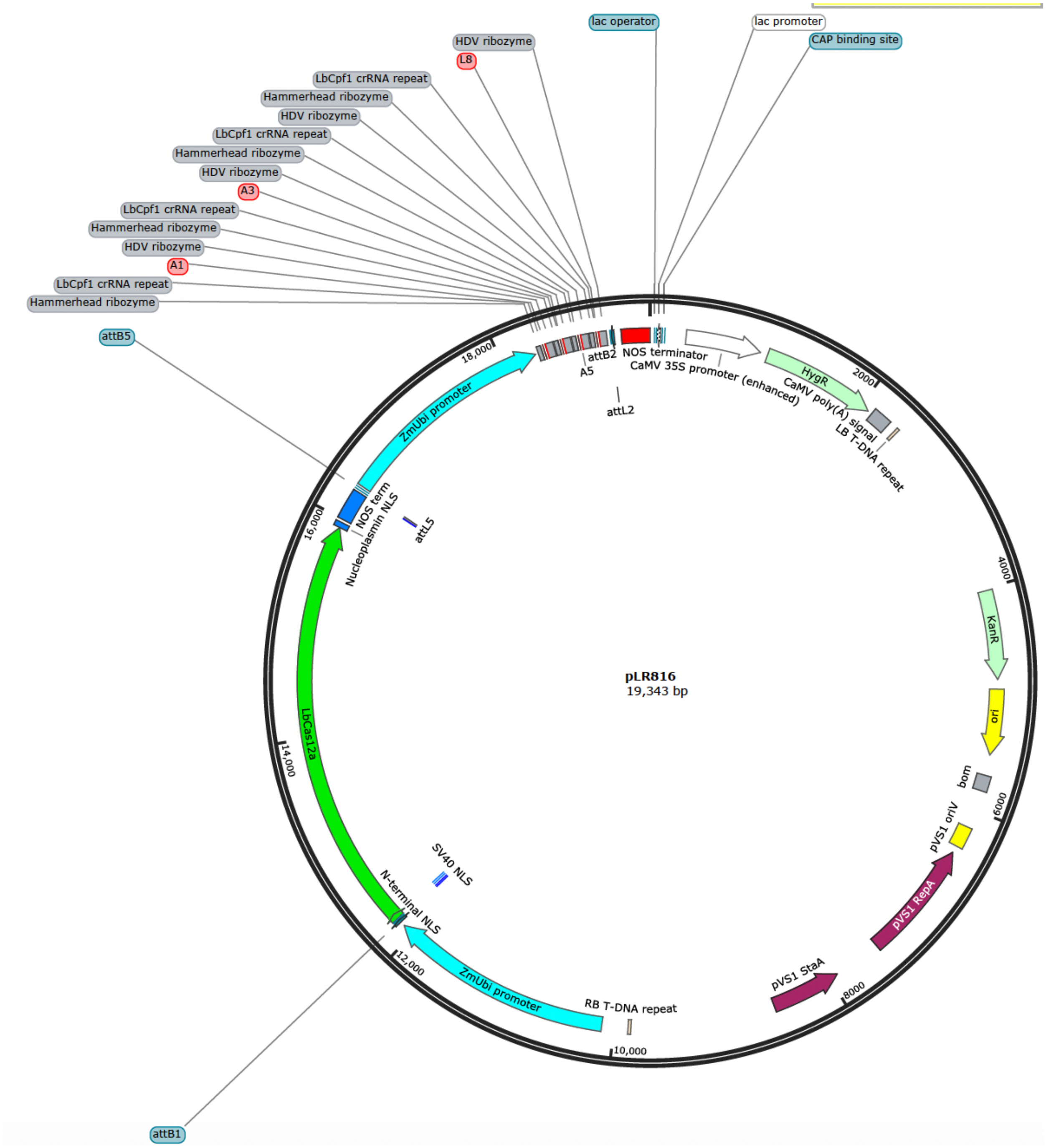


**Supplemental Figure 1. Vector map for LbCas12a based simultaneous expression of four crRNAs in the rice genome.**

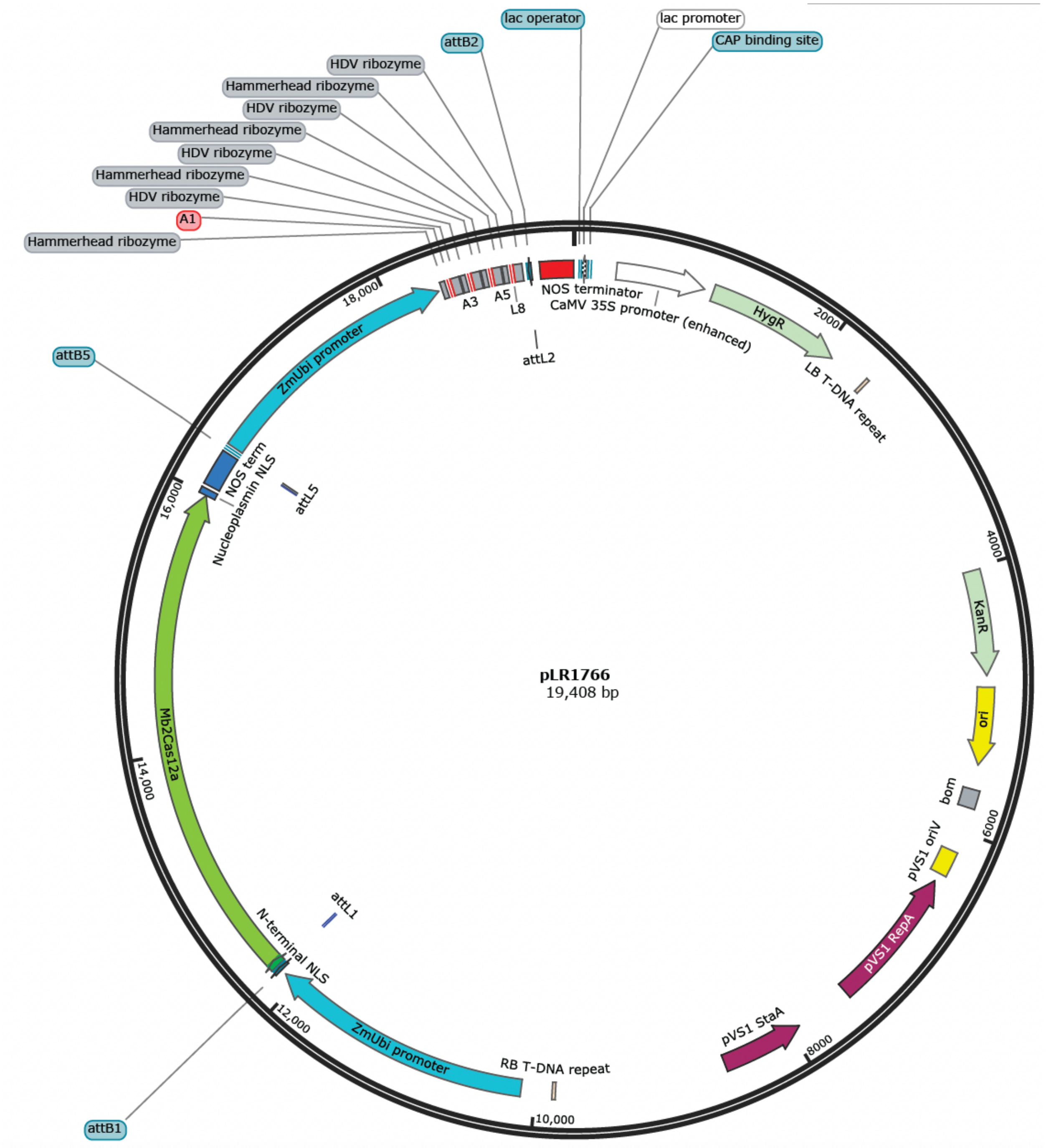


**Supplemental Figure 2. Vector map for Mb2Cas12a based simultaneous expression of four crRNAs in the rice genome.**

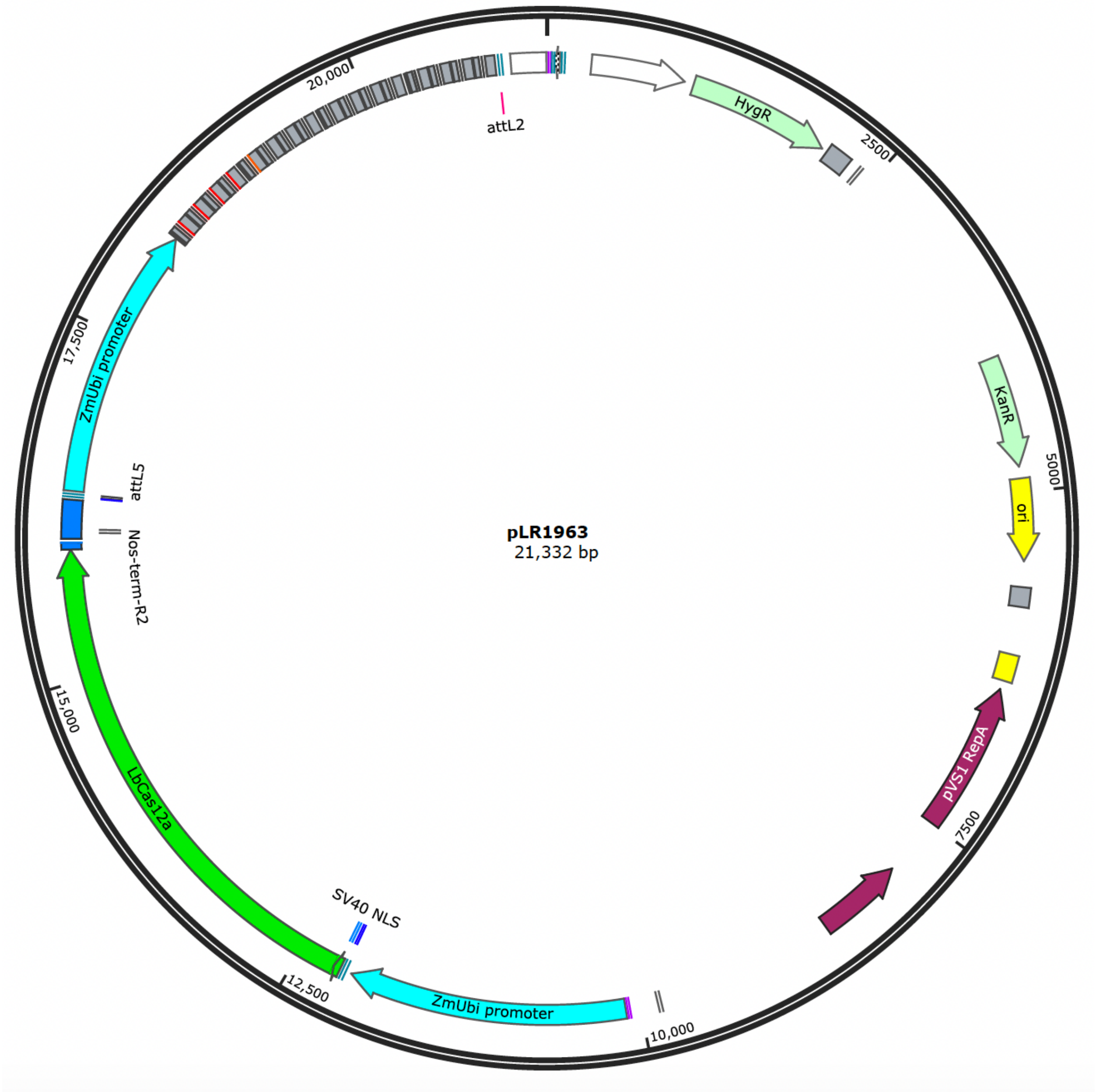


**Supplemental Figure 3. Vector map for LbCas12a based simultaneous expression of 16 crRNAs in the rice genome.**

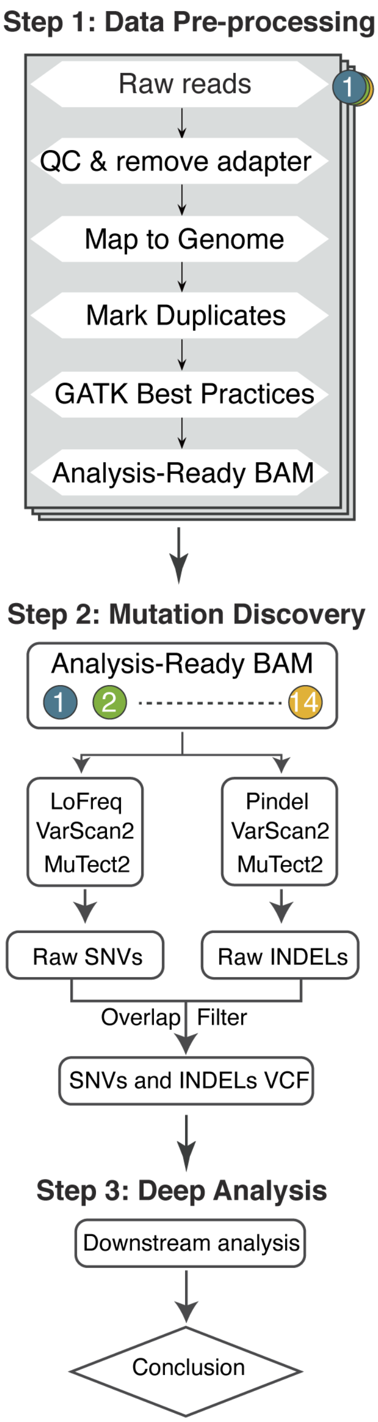


**Supplemental Figure 4. Whole genome sequencing and analysis pipeline.**

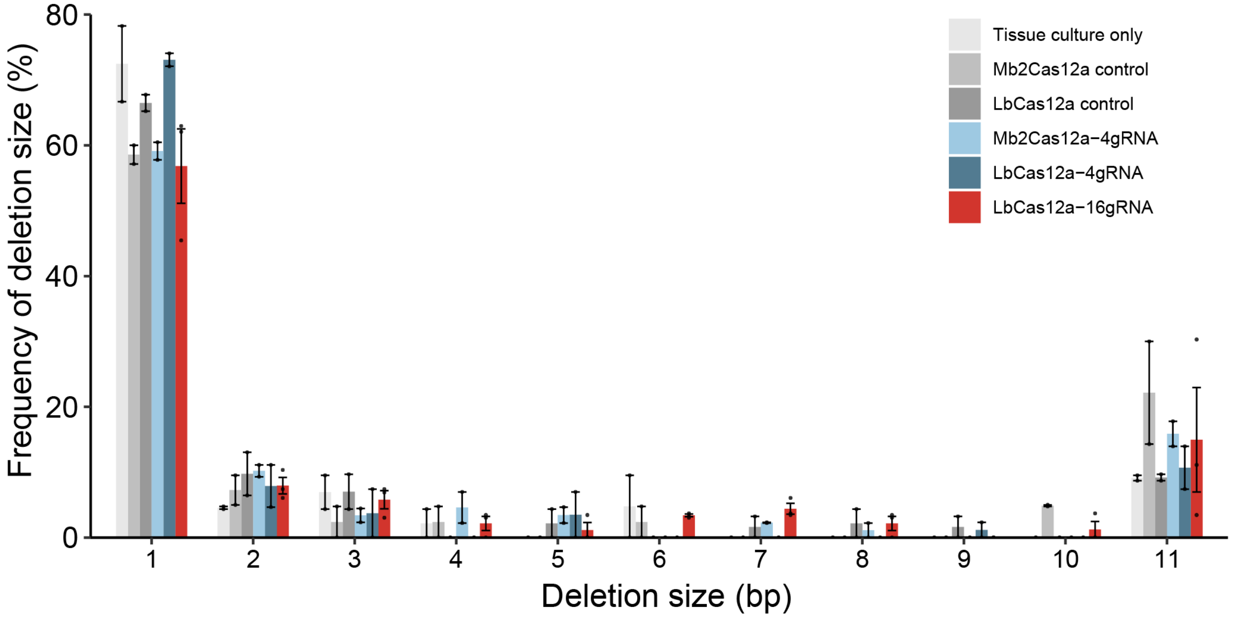


**Supplemental Figure 5. Deletion size analysis of WGS samples.** Error bars represent standard error of the mean (SEM) of two or three lines.

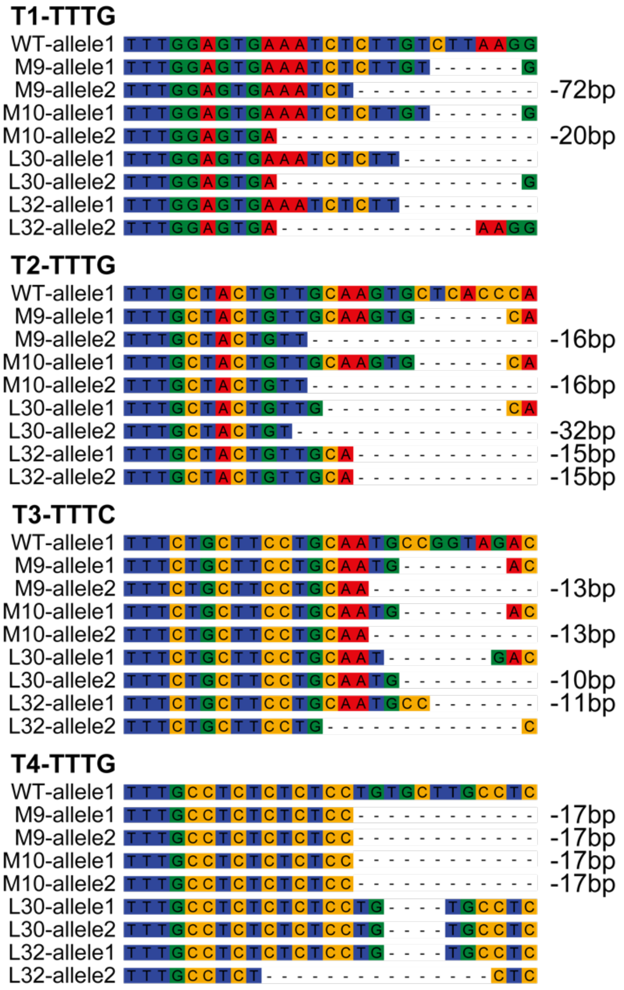


**Supplemental Figure 6. Genotypes of T0 lines with multiplexing four crRNAs by Mb2Cas12a and LbCas12a.** M9 and M10 are two T0 lines edited by Mb2Cas12a. L30 and L32 are two T0 lines edited by Lb2Cas12a. All the lines carried biallelic deletion mutations.

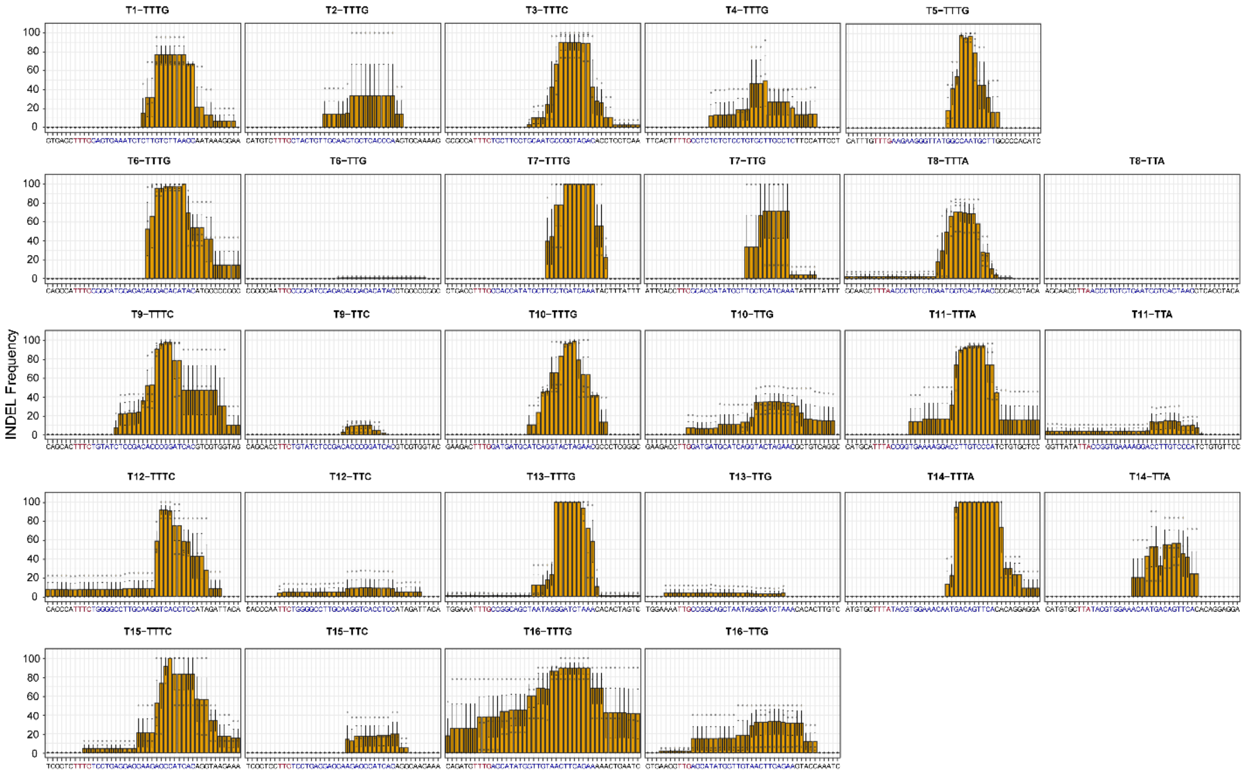


**Supplemental Figure 7. Deletion position at different target sites in T0 lines by LbCas12a with multiplexing 16 crRNAs.** The frequency for each nucleotide position is calculated using the number of reads with this position edited divided by the number of edited reads for this target site. Error bars represent standard error of the mean (SEM) of three lines.

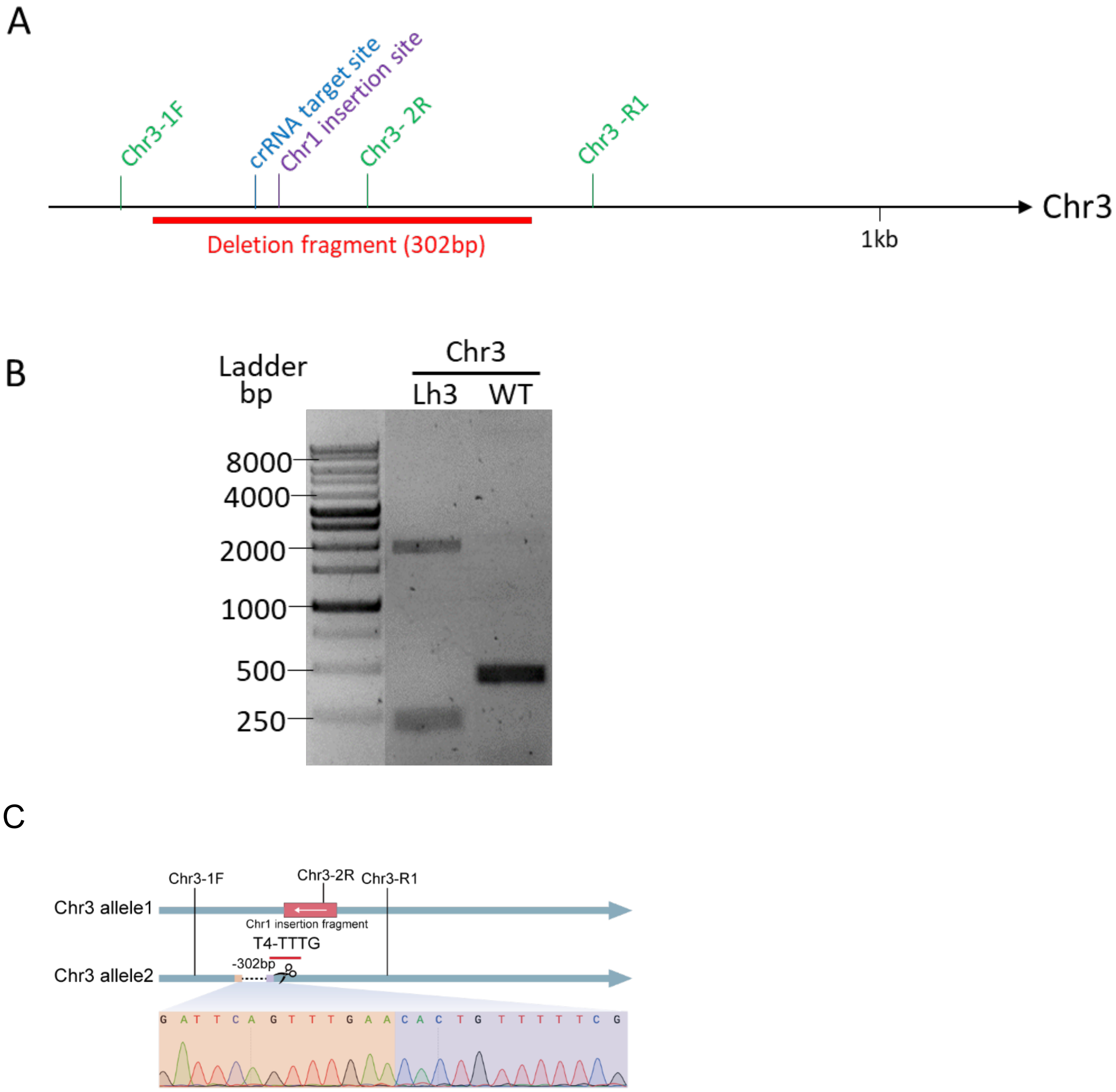


**Supplemental Figure 8. PCR validation of a large DNA insertion and deletion on Chromosome 3 in Lh3.** (A) Location of crRNA target site, insertion site, deletion fragment and PCR primers on Chromosome 3. (B) Gel electrophoresis of PCR product using primers Chr3-1F and Chr3-R1 in Lh3 and WT. (C) Validation of 302 bp deletion on Chromosome 3 in Lh3 by Sanger sequencing.

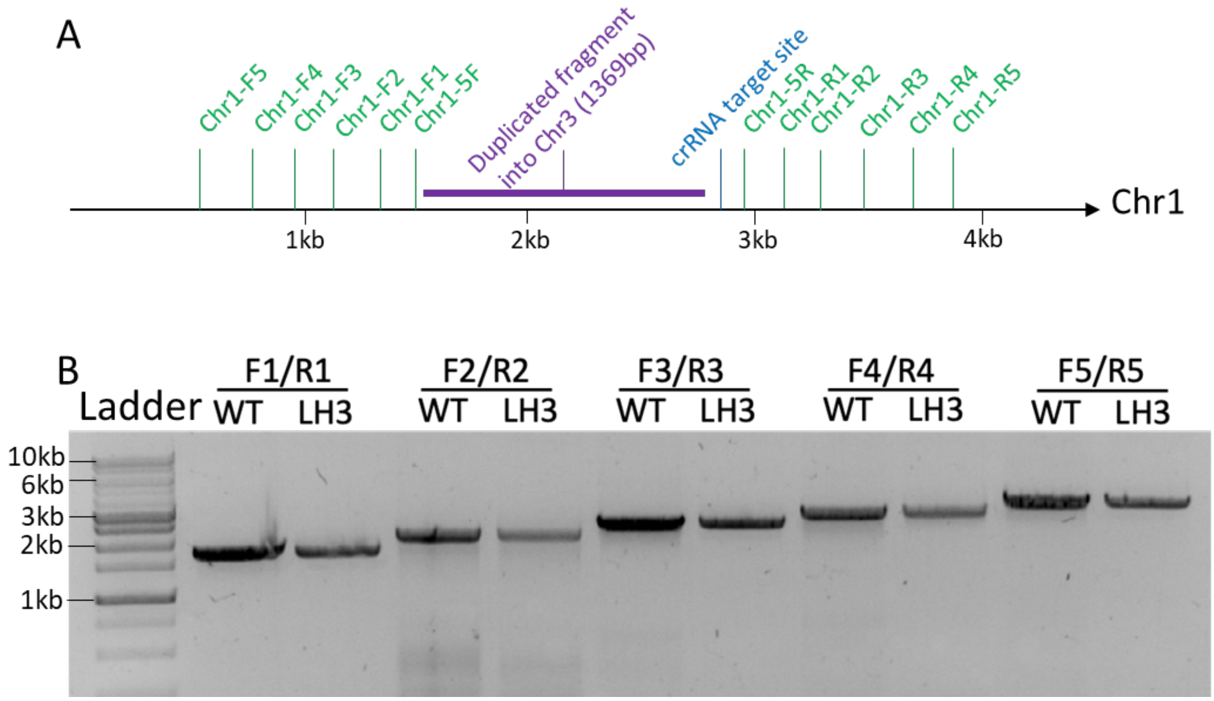


**Supplemental Figure 9. Duplicated DNA sequence in Chromosome 3 is not deleted in Chromosome 1.** (A) Location of crRNA target site, duplicated fragment and PCR primers on Chromosome 1. (B) Gel electrophoresis of PCR products using five pairs of primers in Lh3 and WT.

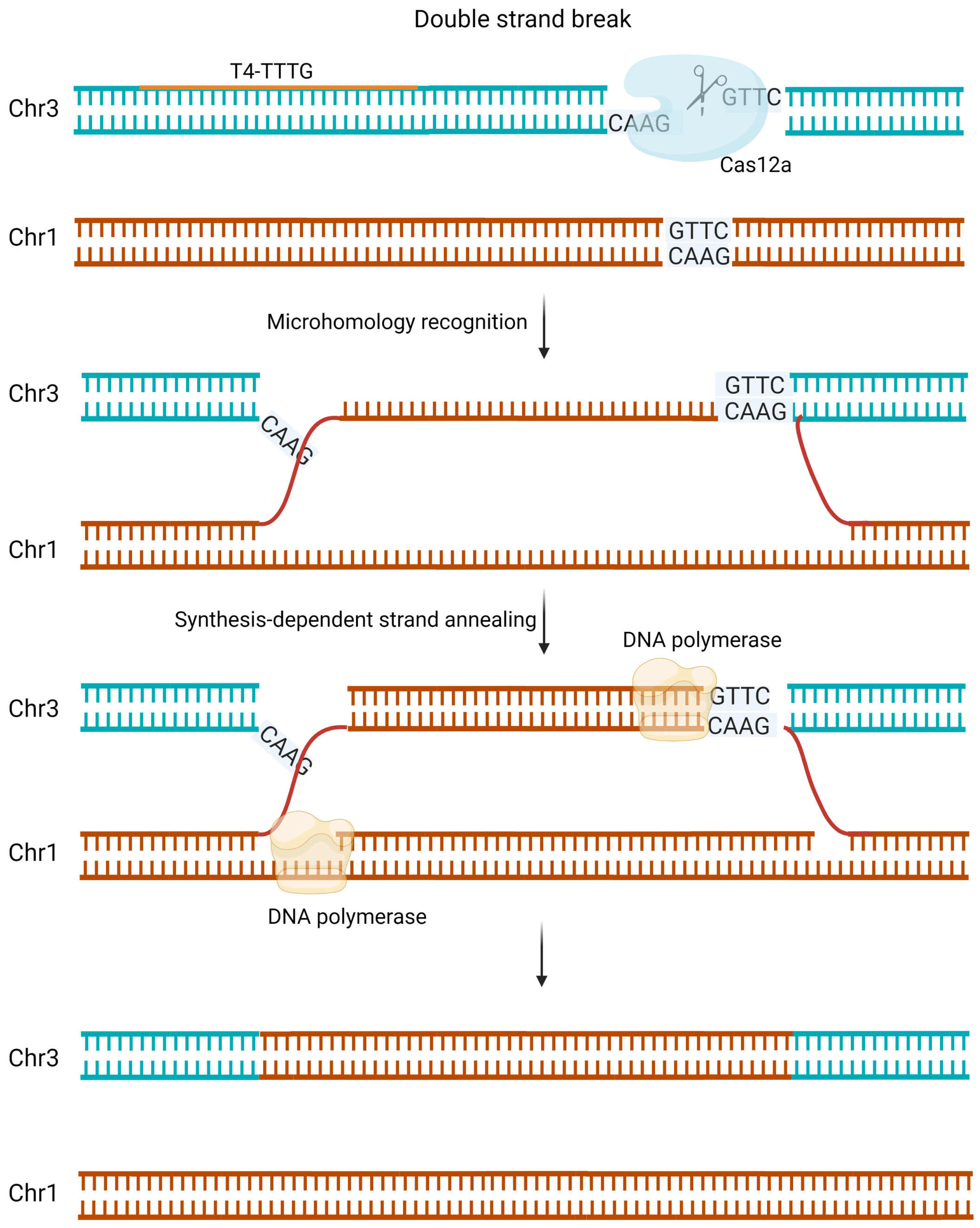


**Supplemental Figure 10. Possible microhomology-mediated synthesis-dependent strand annealing involved in duplication of Chr1 fragment in Chr3.**
